# Bacteriocins in archaea & archaeocins in bacteria

**DOI:** 10.64898/2026.07.27.741052

**Authors:** Romain Strock, Tobias Warnecke

## Abstract

Archaea and bacteria routinely live side by side in microbial communities and must interact at least on occasion. Whether such cross-Domain interactions are dominated by mutual disregard, co-operation, or conflict remains fundamentally unknown. One potential window into archaeal-bacterial conflict is to ask whether some of the molecular weapons bacteria wield to kill other bacteria are present in archaea, and vice versa. Here, to start to address this question, we carry out a phylogenomic survey of bacteriocins in archaeal genomes and archaeocins in bacterial genomes. We find that more than 20% of known bacteriocins – proteins deployed by bacteria against other bacteria - have at least one homolog in archaea. Typically, these archaeal homologs are related to bacteriocins targeting (and encoded by) monoderm bacteria. Based on conservation of functionally critical residues, protein structure, and accessory genes critical for bacteriocin biosynthesis, we highlight homologs of subtilosin A, encoded in some *Thermococcus* archaea, as promising candidates for experimental follow-up work. We also show that halocin C8, originally described in *Natrinema* archaea, is comparatively common in bacterial genomes, including a number of skin-resident *Staphylococcus* species. Our results suggest that bacteriocins/archaeocins are shared across Domain boundaries with some regularity. While many instances are phylogenetically isolated – raising doubts about their functional importance and integration into host physiology – some bacteriocins are present in multiple related genomes and embedded in broader biosynthetic gene clusters that are also found in the original producers, suggesting that archaea and bacteria periodically use the same weapon systems in conflicts with other microbes. Further study of these systems might elucidate cross-Domain conflict and the nature of archaeal-bacterial interactions in different environments.

## INTRODUCTION

In many environments, archaea and bacteria live alongside each other, but the manner of their co-existence – whether they compete with each other, support each other, or largely live parallel lives – typically remains poorly understood (Moissl-Eichinger et al. 2018; Borrel et al. 2020; Strock et al. 2025). In particular, while conflict is common amongst bacteria and amongst archaea (Torreblanca et al. 1994; Shand and Leyva 2007; Atanasova et al. 2013; Besse et al. 2015; Granato et al. 2019; Palmer and Foster 2022; Zachs et al. 2024), we do not know how frequently archaea antagonize bacteria and vice versa, be that through competition for the same limited resources or through more targeted interference that results in growth arrest or even death. In a handful of instances, it has been demonstrated experimentally that exposure to archaeal supernatant can kill bacteria (Shand and Leyva 2008; Atanasova et al. 2013; Megaw et al. 2019; Castro et al. 2021; Liang et al. 2023; Strock et al. 2025; Taissir et al. 2026), but, in almost all cases, the molecular effectors involved remain unknown, let alone how they are deployed and whether they have evolved to target members of the other Domain of Life specifically.

One strategy to identify potential molecular mediators of cross-Domain conflict is to scour archaeal genomes for weaponry already in use as tools of conflict amongst bacteria (Makarova et al. 2019; Strock et al. 2025). Finding homologs of bacterial weaponry in archaeal genomes does not by itself establish that the homolog is used against bacteria, or used antagonistically at all, but it may provide a valuable first lead. We recently illustrated this approach for peptidoglycan hydrolases (PGHs): normally involved in cell wall remodelling during bacterial cell division, prior work had uncovered several instances where PGHs had evolved into weapons, secreted by one bacterium to target and compromise the cell wall of another (Wu et al. 2003; Akesson et al. 2007). Surprisingly, we discovered numerous PGH homologs in archaea – organisms that do not make bacterial peptidoglycan (Strock et al. 2025). We showed that two of these homologs, Woldo and Danwoldo, secreted by the halophilic archaeaon *Halogranum salarium* B-1, kill the halophilic bacterium *Halalkalibacterium halodurans*.

At the time, we pointed out that peptidoglycan hydrolases are likely not unique as molecular mediators of cross-Domain antagonism. In particular, we noted, as have others before us (Makarova et al. 2019), that some archaea encode homologs of a class of proteins collectively known as bacteriocins.

Bacteriocins are genome-encoded and ribosomally synthesized proteins with bactericidal or bacteriostatic activity (Simons et al. 2020). They often undergo (several) enzymatic modifications to yield the final, biologically active molecule and are usually embedded in biosynthetic gene clusters (BGCs), flanked by genes involved in their regulation, post-translational processing, transport, and/or secretion. Whenever bacteria target closely related strains (Akesson et al. 2007; Azevedo et al. 2015) – which they often do, as these strains are prime competitors for the same niche – they run the risk of succumbing to their own toxins. As a consequence, bacteriocin BGCs often also include immunity proteins that nullify these effects, for example by binding to and neutralizing the bacteriocin.

Like many other genes involved in interbacterial conflict (Kogay et al. 2024; Granato et al. 2025), bacteriocins are often transferred horizontally to other bacteria (Rossi et al. 2014; Krauss et al. 2022). But have bacteriocins, and the broader gene clusters required for their biosynthesis and deployment, also been shared with archaea? If so, have these genes been functionally integrated into archaeal genomes? Have they retained their original function and are secreted to kill bacteria?

Here, as a first step towards resolving these questions, we present an up-to-date census of known bacteriocins in archaeal genomes. Conversely, we survey archaeocins – proteins secreted by archaea that kill other archaea (O’Connor and Shand 2002; Besse et al. 2015) – in bacterial genomes. Considering phylogenetic spread, BGC completeness, structural conservation and other features, we highlight one bacteriocin (subtilosin A) and one archaeocin (halocin C8), that have made their way into genomes from the other Domain of Life, that likely still act as tools of conflict, and that are therefore promising candidates for future experimental investigation.

## RESULTS

### Surveying archaeal genomes for homologs of known bacteriocins

The BAGEL4 database (van Heel et al., 2018) compiles protein sequences of known bacteriocins, including unmodified as well as ribosomally synthesized and post-translationally modified peptides (RiPPs). Extending our earlier preliminary work (Strock et al. 2025), we used the core peptide database of BAGEL4 to seed searches against a phylogenetically balanced subset of the Genome Taxonomy Database (GTDB, see Methods), which contains 3,706 archaeal and 50,640 bacterial genomes.

Surveying archaeal genomes in our database, we find putative homologs for 85 out of 413 de- duplicated bacteriocins (21%), spread across 327 archaea (Table S1), equivalent to 8% of archaea in the database. 58 bacteriocins are only found in a single archaeal genome, raising doubts about their functional integration and adaptive benefit to the new host genome; these instances might simply represent short-lived HGT events that are inconsequential for fitness. As our prime motivation is to identify promising systems for future experimental investigation, we narrowed our survey to bacteriocins that have been detected in at least two archaeal genomes. To further focus on the most promising candidates, we reviewed the primary literature on these 27 bacteriocins and only considered cases where antibacterial activity has been demonstrated experimentally (see Methods), leaving 15 distinct bacteriocins (Figure 1).

**Figure 1.**
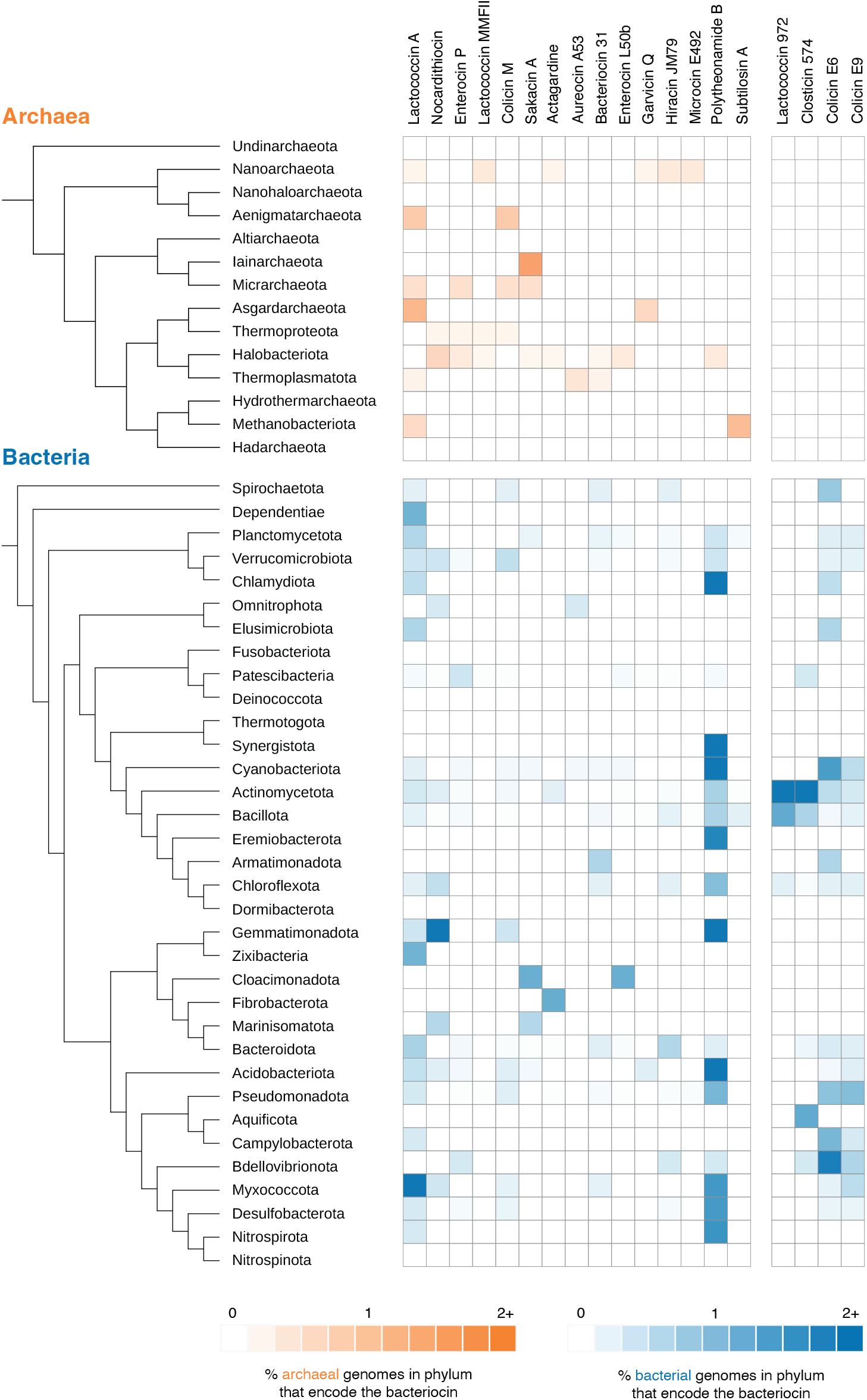
Relative incidence of known bacteriocins across archaeal and bacterial phyla. The 15 bacteriocins highlighted in the main text are shown in the left-hand block. The right-hand block illustrates, for a selection of well-known examples, that not all bacteriocins have homologs in archaea.

Eleven of the 15 bacteriocins (73%) are produced by and target monoderm bacteria. With approximately twice as many diderm bacteria in our database, the odds of drawing eleven monoderm bacteria by chance is about 1 in 2,000 (one-sided binomial test, P= 4.7e-4 < 0.05). *Listeria* species are common targets (see below), but – without systematic testing of antibacterial activity across a large, unbiased panel of species – it is not possible to rule out that this is simply the result of ascertainment bias. Eight of the 15 bacteriocins (53%) are from lactic acid bacteria belonging to *Enterococcus* (N=4), *Lactococcus* (N=3) or *Lactobacillus* (N=1) species. Amongst bacteriocins for which the mechanism of action is known, membrane disruption appears to be common. Below, we will provide a brief overview of the 15 bacteriocins with ≥2 homologs in archaea and then move on to examine one of them, subtilosin A, in greater detail.

### Bacteriocins from Lactococci

The most common archaea-encoded bacteriocins in our set (N=8 instances in archaea, Figure 1) are homologs of Lactococcin A, a well-characterised protein produced by *Lactococcus lactis* subsp. *cremoris*, which kills related Lactococci strains by binding to the mannose phosphotransferase system (man-PTS), thereby forming pores in the cytoplasmic membrane (Holo et al. 1991; Li et al. 2023). Garvicin Q (N=2 instances), produced by *Lactococcus garvieae* BCC 43578, also targets the man-PTS system, and is active against different *Listeria* species. Unlike Lactococcin A, Garvicin Q does not appear to target other *Lactococus* species (Tymoszewska et al. 2017). A third Lactococcus-derived bacteriocin, Lactococcin MMFII (N=4 instances), is again active against *Listeria* species. Its mechanism of action is unknown (Ferchichi et al. 2001). Archaeal homologs of these bacteriocins are found in Nanoarchaeota, Asgardarchaeota, and isolated genomes in other clades, including *Methanobacterium bryantii,* which encodes a lactococcin A homolog (Figure 1, Table S1).

### Bacteriocins from Enterococci

Four archaea-encoded bacteriocins were originally described in Enterococci: Enterocin P (N=4 homologs in archaea), Enterocin L50b (N=2), Hiracin JM79 (N=2), and bacteriocin 31 (N=2). Enterocin P is produced by *Enterococcus faecium* T136 and forms pores in the membrane of sensitive bacteria, including, once again, *Listeria monocytogenes* (Herranz et al. 2001). Bacteriocin 31, produced by *Enterococcus faecalis* YI717, is active against *Enterococcus hirae*, *E. faecium*, and *L. monocytogenes* (Tomita et al. 1996). Enterocin L50b is produced by *E. faecium* L50 and targets several Gram-positive bacteria, including *Lactococcus lactis* (Cintas et al. 1998). Hiracin JM79 (N=2), produced by *E. hirae* DCH5, is also active against Gram-positive strains, including *Enterococcus*, *Lactococcus* and *Listeria* species (Sánchez et al. 2007). Five of the ten archaeal homologs of enterococcal bacteriocins are found in members of the Halobacteriota.

### Bacteriocins from other Gram-positive bacteria

Sakacin A (N=3 instances in archaeal genomes) from *Lactobacillus sakei* DSMZ 6333, another bacteriocin that is active against *Listeria*, acts by interfering with transmembrane potential and disrupting the pH balance of susceptible cells (Trinetta et al. 2012).

Nocardithiocin (N=5) was isolated from *Nocardia pseudobrasiliensis* IFM 0757 (phylum Actinomycetota), a soil dweller and opportunistic human pathogen. It has activity against a range of Gram-positive bacteria, most notably against rifampicin-resistant strains of *Mycobacterium tuberculosis,* but also against related *Nocardia* strains (Mukai et al. 2009). Interestingly, four out of the five archaeal hits are from *Methanosarcina* species (including *Methanosarcina mazei*), meaning that 40% (4 out of 10) of the *Methanosarcina* genomes in our database encode homologs of Nocardithiocin.

Actagardine (N=2) is produced by *Actinoplanes garbadinensis* ATCC31049 and disrupts cell wall synthesis of Gram-positive bacteria including *E. faecalis*, *Staphylococcus aureus* and *Streptococcus pneumoniae* (Boakes et al. 2009).

Aureocin A53, found in two Thermoplasmatota genomes, is produced by the opportunistic pathogen *S. aureus* A53 and has been shown to kill related *Staphylococcus* strains by dissipating membrane potential (Netz et al. 2002).

Subtilosin A is produced by *Bacillus subtilis* str. 168 and acts by permeabilizing the cytoplasmic membrane of susceptible bacteria. It exhibits a broad range of activity against Gram-positive and Gram-negative bacteria, including against *Klebsiella pneumoniae*, *Streptococcus pyogenes* and *L. monocytogenes* (Babasaki et al. 1985; Shelburne et al. 2007). Homologs of subtilosin A, to which we will return below, are found in two archaea in our database: *Thermococcus celer* and *Thermococcus nautili*.

### Bacteriocins from Gram-negative bacteria

Colicin M (N=3), produced by several *E. coli* strains, disrupts peptidoglycan synthesis of susceptible *E. coli* competitors (Chérier et al. 2021). Microcin E492 (N=2), found in two members of the Nanoarchaeota, is a short peptide from *Klebsiella pneumoniae* that depolarises the membrane of sensitive strains, including *E. coli* K-12 (Lorenzo and Pugsley 1985) .

Finally, polytheonamide B (N=2), originally thought to be a non-ribosomal peptide due to its highly modified nature, has in fact been shown to originate from a precursor peptide that goes through 48 post-translational modifications. It is produced by an uncultured bacterial marine sponge symbiont and depolarises the membrane potential of susceptible Gram-positive bacteria (Freeman et al., 2012). In archaea, it is found in two species of the genus *Natronorubrum*.

### Subtilosin A in Thermococcales archaea

As highlighted above, finding a homolog does not guarantee that this homolog acts in the same manner as the original bacteriocin. The archaeal homolog might no longer be active against bacteria. It might, instead, be active against archaea, or eukaryotes. It might even have undergone a drastic change in function and no longer be involved in microbial conflict at all. Or it could be a transient passenger that has recently arrived via horizontal gene transfer and might be capable of antibacterial action in principle but has not been functionally integrated into the archaeal genome, and perhaps never will be.

To identify promising candidates for future experimental studies, we therefore examined features that suggest continued deployment as a bacteriocin: sequence and structural homology, conservation of accessory genes, and phylogenetic spread in archaeal genomes. We also considered whether the homolog was found in high-quality, complete genomes, which would increase one’s confidence that a) the bacteriocin is assigned to the right organism rather than being an assembly contaminant and that b) we can fully capture any associated BGC. Finally, we considered whether the mechanism of action and pathway of biogenesis are understood for the original bacteriocin, which would facilitate downstream testing of the archaeal homolog.

Jointly considering these criteria, we decided to focus on archaeal homologs of subtilosin A. Subtilosin A is a member of the sactipeptide family of proteins, which feature characteristic sactionine rings, where <u>s</u>ulphur to <u>a</u>lpha <u>c</u>arbon thioether (‘sacti’) cross-links connect a cysteine residue and the alpha carbon of another amino acid (Chen et al. 2021). Subtilosin A is notable for its broad spectrum of activity against many Gram-positive and Gram-negative bacteria, including several human pathogens (Shelburne et al. 2007). It also exhibits spermicidal and antiviral properties (Sutyak et al. 2008; Quintana et al. 2014). Activity against archaea has, to our knowledge, not been reported.

The precursor of subtilosin A (sboA) in *B. subtilis* 168 is a 43-amino acid peptide encoded as part of the *sbo-alb* operon (Figure 2A). The so-called ‘<u>a</u>nti-listerial <u>b</u>acteriocin’ (*alb*) locus is made up of seven genes involved in maturation, transport and immunity: AlbA is a radical S-adenosylmethionine (SAM) enzyme that catalyses the formation of three key thioether cross- links in sboA (Flühe et al. 2012) (Figure 2B). AlbE and AlbF form a complex responsible for cleaving the leader peptide and closing the ring to form the mature cyclic subtilosin A molecule (Ishida et al. 2022) (Figure 2B). AlbC is an ABC transporter protein primarily responsible for the export of the mature peptide. Mutational analysis has shown that AlbA, AlbC and AlbF are essential for the production of subtilosin A, whereas AlbB, AlbC and AlbD are necessary for appropriate immunisation – through an unknown mechanism – of the producer against its own bacteriocin (Zheng et al. 2000). The function of AlbG is unknown but its deletion leads to a reduced amounts of active peptides in the supernatant (Zheng et al. 2000).

**Figure 2.**
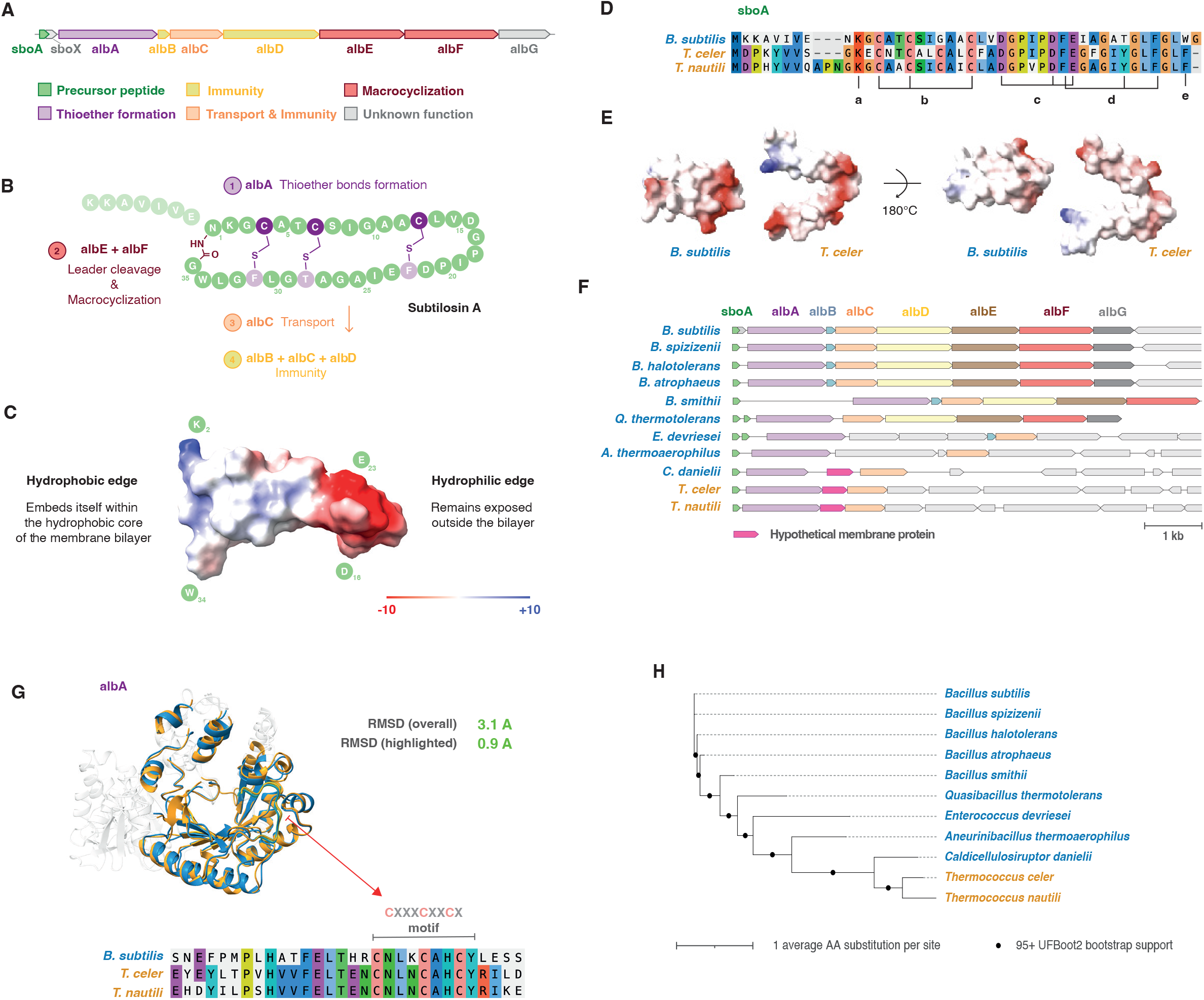
Subtilosin A and its archaeal homologs. **A.** Schematic representation of the subtilosin A BGC in *B. subtilis*. **B.** Ilustration of subtilosin A processing, highlighting the relevant BGC members involved. **C.** Structural representation of subtilosin A, highlighting its amphipatic nature. **D.** Alignment of subtilosin A and its homologs in *T. celer* and *T. nautilii*, indicating the lysine (a) and tryptophan (e) involved in membrane permeabilization, three cysteines (b) and two phenylalanines (d) involved in thioether bond formation, and three conserved negatively charged amino acids forming the hydrophilic edge (c). **E.** The *T. celer* homolog of subtilosin A has a conserved amphipatic nature. **F.** Gene neighbourhood of subtilosin A homologs. **G.** Structural and sequence alignment of AlbA, highlighting the conserved CXXXCXXCX motif. **H.** Protein tree of a concatenated alignment of sboA, AlbA, and AlbC.

A peculiar feature of the subtilosin A gene cluster is the presence of a second bacteriocin-like precursor gene, *sboX*, whose coding sequence overlaps with that of *sboA* (**Error! Reference source not found.**A). The product of *sboX* resembles a precursor of type II bacteriocins and is expressed in stationary phase anaerobic cultures of *B. subtilis*. Its function is unknown, but an in-frame deletion has no impact on the activity, production of, or immunity to subtilosin A (Zheng et al. 2000), demonstrating that sboX is dispensable for minimal subtilosin A function.

Subtilosin A is amphipathic (Figure 2C) and this property plays a key role in its activity. The hydrophobic edge of the peptide partially inserts itself into the membrane core, while its hydrophilic edge remains exposed, ultimately leading to membrane permeabilization (Thennarasu et al. 2005).

Our initial search, described above, revealed homologs encoded by two hyperthermophilic archaea: *Thermococcus celer* (strain Vu 13) and *Thermococcus nautili* (strain 30-1). Both species encode the subtilosin A precursor gene *sboA* on their main chromosome.

Alignment of *B. subtilis* sboA with its Thermococcus homologs highlights conservation of key amino acids important for subtilosin A activity: the three cysteines involved in thioether bond formation in *B. subtilis* sboA are conserved, as are the two phenylalanine residues that participate in the outer thioether bonds (Figure 2D). The central threonine of the final thioether bond in *B. subtilis* is a tyrosine in the archaeal homologs (Figure 2D). The amphipathic nature of subtilosin A also appears conserved (Figure 2E). A tryptophan residue on the hydrophobic edge of subtilosin A, one of two residues shown to be involved in the membrane permeabilization activity (Thennarasu et al., 2005), has been swapped for phenylalanine, another hydrophobic amino acid; the other, a lysine, is conserved.

The tertiary structures of sboA and sboA-like precursor proteins from *B. subtilis* and *T. celer,* the former a crystal structure, the latter predicted with AlphaFold 3 (Abramson et al. 2024), have a similar charge distribution. The *T. celer* homolog displays a more open conformation (Figure 2E), but one might expect that both conformations exist, with a closed conformation enforced by the addition of thioether bonds between the N- and C-terminal halves of the mature protein.

Examining the genomic neighbourhood of *sboA*, we find that two other genes found in the *B. subtilis* BGC are present in all Thermococcus genomes that encode *sboA*, but absent from all genomes that do not: *albA* and *albC* (Figure 2F).

A CXXXCXXCX motif in AlbA, characteristic of radical SAM enzymes, is essential for its ability to catalyze thioether bond formation (Flühe et al. 2012). This motif is conserved in *T. celer* and *T. nautili* and the fold surrounding the motif shows good structural homology to AlbA (Figure 2G). It therefore seems reasonable to assume that AlbA interacts with sboA in a similar manner in *B. subtilis* and the Thermococcales species examined here.

Two proteins involved in immunity in *B. subtilis*, AlbB and AlbD, are missing, as are AlbE and AlbF, the proteins responsible for cleaving the leader peptide and closing the sactionine ring, and the two genes of unknown function, AlbG and sboX. For proteins solely involved in immunity (AlbB and AlbD, but not AlbC, which has an additional transport function), loss in the archaea might not be unexpected: if Thermococcus sboA continues to target bacteria, archaea – thanks to their divergent membrane and cell wall composition – might be naturally immune. Regarding AlbE and AlbF, it is interesting to note that processing intermediates of sboA prior to macrocyclization still exhibit antibacterial activity, albeit reduced (Jia et al. 2025), suggesting that AlbE and AlbF are, in fact, not strictly required for antibacterial function.

To better understand the evolution of the sboA BGC in the Thermococcales and how they might have acquired it, we surveyed bacterial genomes in our database for those that harbour the minimal triumvirate of *sboA*, *albA* and *albC*. This search yielded eleven genomes (including the original producer *B. subtilis*), exclusively from the phylum Bacillota, four of which are thermophiles: *Bacillus smithii, Quasibacillus thermotolerans*, *Aneurinibacillus thermoaerophilus* and *Caldicellulosiruptor danielii*. A protein tree built from the concatenated alignment of sboA, AlbA and AlbC (Figure 2H) shows Thermococcales branching with *C. danielii*, which was isolated from a hot spring in Japan and grows optimally at 75°C (Manesh et al. 2024). Interestingly, *C. danielii* also lacks AlbB, D, E, F and G but encodes a protein of unknown function in between its *albA* and *albC* homologs (Figure 2F). A structurally very similar protein, which exhibits transmembrane features, is also found in the sboA-encoding Thermococcales genomes and absent from those not encoding sboA (Figure 2F).

These findings are consistent with an evolutionary scenario where a minimal subtilosin-like pathway evolved in bacteria and was subsequently acquired via horizontal transfer by a Thermococcus archaeon from a *Caldicellulosiruptor*-related bacterium that inhabited the same high-temperature niche. Whether this homolog of sboA has retained its antibacterial activity in either *C. danielii* or any of the Thermococcus species will need to be confirmed experimentally.

### Archaeocins in bacteria

Archaea too engage in conflicts amongst each other, including through the use of proteins that act in a manner analogous to bacteriocins and are therefore known as archaeocins (Torreblanca et al. 1994; O’Connor and Shand 2002; Atanasova et al. 2013). In stark contrast to bacteria, only a very small number of such proteins have been described in detail and the scale and pervasiveness of negative interactions involving different archaea remains fundamentally unknown. Almost all known tools involved in inter-archaeal conflict come from halophilic archaea where they are known as halocins (O’Connor and Shand 2002; Besse et al. 2015). Just like their bacterial counterparts, these halocins are genome-encoded, often target related organisms, and therefore require immunity mechanisms to protect the producer.

Given that archaea encode bacteriocins, it is pertinent to ask whether the inverse is also true: Do bacteria encode homologs of archaeocins and, if they do, are these deployed against archaea?

Three families of halocins have been described in molecular detail: H4, C8 and S8. The latter two belong to the same protein superfamily (Makarova et al. 2019). Other named halocins have been identified biochemically, for instance halocins H1, SH10 and Sech7a (Platas et al. 2002; Pašić et al. 2008; Karthikeyan et al. 2013), but their amino acid and precursor gene sequences are not known. We therefore only consider halocins H4, C8 and S8, but expand our search to the entirety of GTDB release 214.

Makarova and colleagues previously noted the presence of C8 homologs in a number of bacteria, predominantly Firmicutes/Bacillota (Makarova et al. 2019). We recapitulate these findings, recovering a comparatively large number of C8 homologs in bacteria (N=26) and no S8 homologs (Table S2). In contrast to this earlier work, we do find a small number of H4 homologs (N=4).

### Bacterial homologs of halocin H4

Halocin H4 was originally described from *Haloferax mediterranei* R4 (Meseguer and Rodriguez-Valera 1985). Characterization of its hit kinetics indicated that a single protein might be sufficient to cause cell death. Exposure to H4 affects the electrochemical gradients of sensitive archaeal cells (Meseguer and Rodriguez-Valera 1986), and a transmembrane region (at positions 179–194) is essential for archaecidal activity (Chen et al. 2024), suggesting a mode of action that compromises membrane integrity. Halocin H4 has broad-spectrum activity against halophilic organisms, including members of the genera *Haloferax*, *Haloarcula,* and *Halorubrum*. Importantly, H4 was recently shown to be active against two bacteria: environmental isolates belonging to the *Aliifodinibius* and *Salicola* species groups (Chen et al. 2024). By implication, bacterial homologs might act on bacteria rather than archaea, so the presence of H4 in a bacterial genome is no conclusive indicator of cross-Domain conflict.

Our survey detects halocin H4 homologs in six species from the archaeal class Halobacteria (from genera *Haloferax*, *Natrinema*, *Natrialba*, *Halomicrobium* and *Haloviva*), but it is not exclusive to halophilic archaea. Other hits include Syntropharchaeia (N=4), Thermococci (N=3), two Methanosarcina species (including *Methanosarcina barkeri*), two metagenome-assembled Thermoplasmata and one Methanomicrobia (*Methanofollis formosanus*).

We found bacterial homologs in three genomes belonging to the genus *Paenibacillus – Paenibacillus amylolyticus*, *Paenibacillus taichungensis*, and *Paenibacillus* sp001955925 – and in the genome of *Natranaerovirga hydrolytica*, a halophilic bacterium growing optimally at 1M NaCl (Sorokin et al. 2012). All of these bacterial H4 homologs carry a signal peptide for secretion via the Sec pathway, different from the Tat-mediated secretion of haloarchaeal H4 proteins.

A phylogenetic tree based on the alignment of halocin H4 homologs across prokaryotes (Figure 3A, see Methods) indicates that all four bacteria form a monophyletic group branching together with the two Thermoplasmata species, whose H4 homologs also harbour Sec signal peptides suggesting that the transition to Sec-mediated secretion preceded the acquisition of H4 by bacteria.

**Figure 3.**
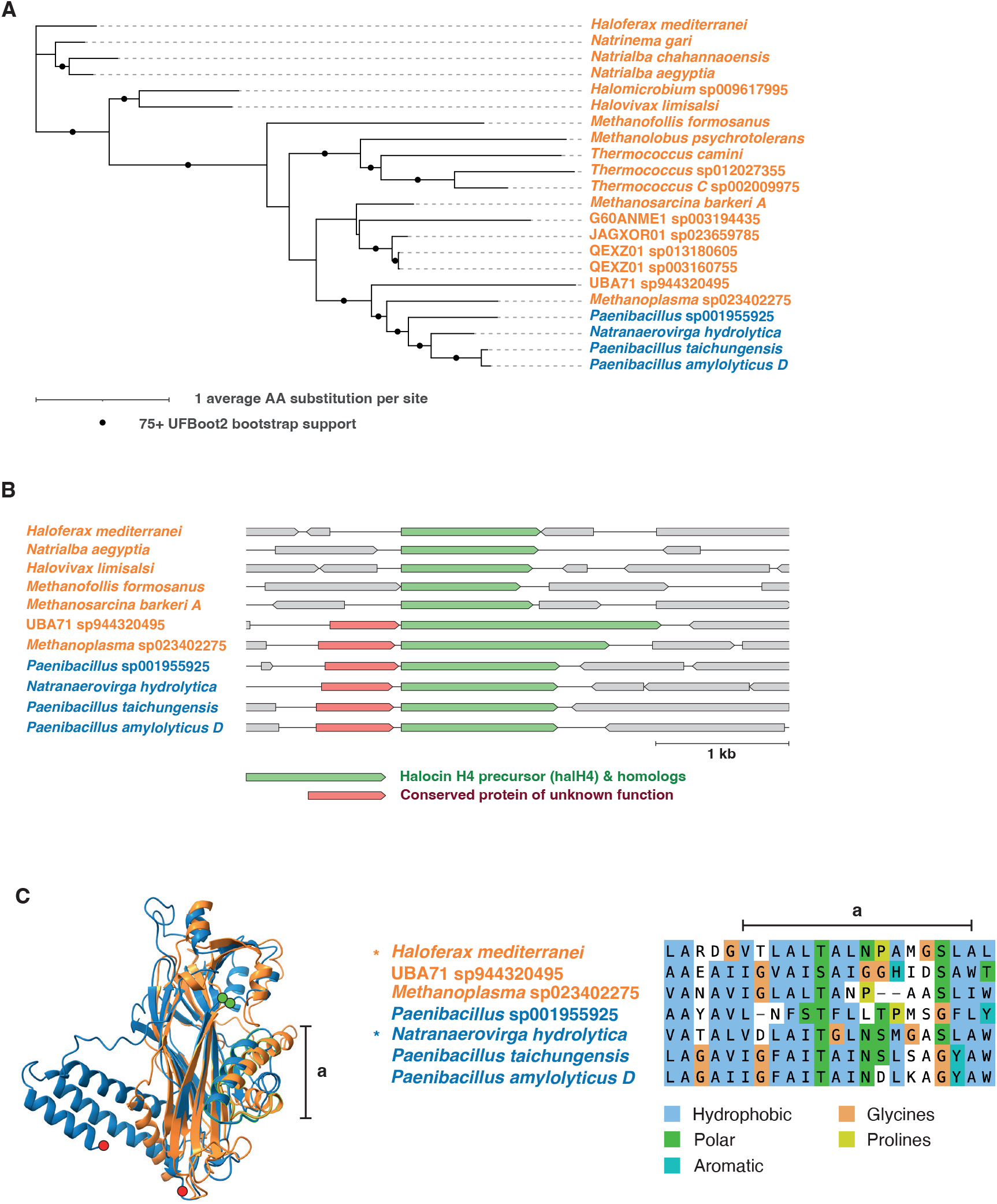
Halocin H4 and its bacterial homologs. **A.** Protein tree of halocin H4 homologs. **B.** Gene neighbourhood of halocin H4 homologs in archaeal and bacterial genomes. **C.** Structural alignment of halocin H4 from *Haloferax mediterranei* and its bacterial homolog in *Natranaerovirga hydrolytica* alongside an excerpt of the protein sequence alignment focusing on the transmembrane region of H4 (a).

Synteny analysis reveals a hypothetical gene upstream of the *halH4* homologs that is conserved in the two Thermoplasmata species as well as all four bacterial species (Figure 3B); its significance remains unknown.

Structurally, the core of H4 seems well conserved between archaea and the four bacteria, as exemplified by the structural alignment between halocin H4 from *H. mediterranei* and its bacterial homolog from *N. hydrolytica* (Figure 3C). The main difference between the two proteins is found at the C-terminus, where the *N. hydrolytica* homolog contains a 58 amino acid extension.

The transmembrane region whose deletion led to loss of activity in *H. mediterranei* (Chen et al. 2024) is poorly predicted by AlphaFold 3 in *H. mediterranei* H4 and its homologs but overall composition in terms of amino acid properties appears well conserved (Figure 3C), suggesting that the membrane-perforating capacity of archaeal H4 may also be conserved in its bacterial homologs.

### Homologs of halocin C8 are common in bacteria

Halocin C8, originally described from *Natrinema* sp. AS7092 (Li et al. 2003), has broad activity against other haloarchaeal strains, including *Halobacterium salinarum* DSM 669, *Haloferax volcanii DS2*, *Natrialba magadii* NCMB2190*, Haloarcula hispanica* and *Halorubrum saccharovorum*. But toxic effects are not universal. For example, *Halobacterium salinarum* DSM 670, *Haloferax denitrificans* and *Natrinema versiforme* appear immune (Li et al. 2003). No activity against bacteria has been reported. Halocin C8 is encoded by the halC8 gene, whose product, ProC8 (UniProt: P83716), is a 283-amino acid protein.

Following secretion via the Tat pathway, this precursor protein undergoes proteolytic cleavage to release an N-terminal immunity protein (HalI) and the 76-amino acid C-terminal mature halocin C8 peptide (Sun et al. 2005). The cleavage mechanism remains unknown.

The *halC8* gene is flanked by genes that may be involved in its regulation, processing and transport. The gene directly upstream of halC8 is a transcriptional regulator dubbed *halR* and the three genes downstream code for ABC transporters (*halT1, halT2* and *halT3*). A gene of unknown function named *halU*, upstream of *halR*, is conserved in several other *Natrinema* and *Haloterrigena* species (Besse et al. 2017).

The mechanism of action of halocin C8 has not been fully elucidated. However, observations using transmission electron microscopy of cells treated with halocin C8 have shown that cells swell and the cell wall at both poles becomes nicked, leading to leakage of cellular content. After 24 hours of treatment, all cells were completely lysed and only cell debris remained, suggesting that C8 activity is targeted against the cell wall (Li et al. 2003)

Consistent with prior results (Makarova et al. 2019), our search identified a fairly large number of C8 homologs in bacteria (N=26), along with 82 homologs in archaea. A maximum likelihood phylogenetic tree built from these 108 homologs suggests that bacteria acquired halocin C8 genes from archaea on at least three separate occasions (Figure 4A). The main group of bacteria (22 of 26 homologs) is composed chiefly of Bacillota bacteria, at least five of which (four *Halobacillus* species and *Alteribacillus bidgolensis*) are known to thrive at high salinity.

**Figure 4.**
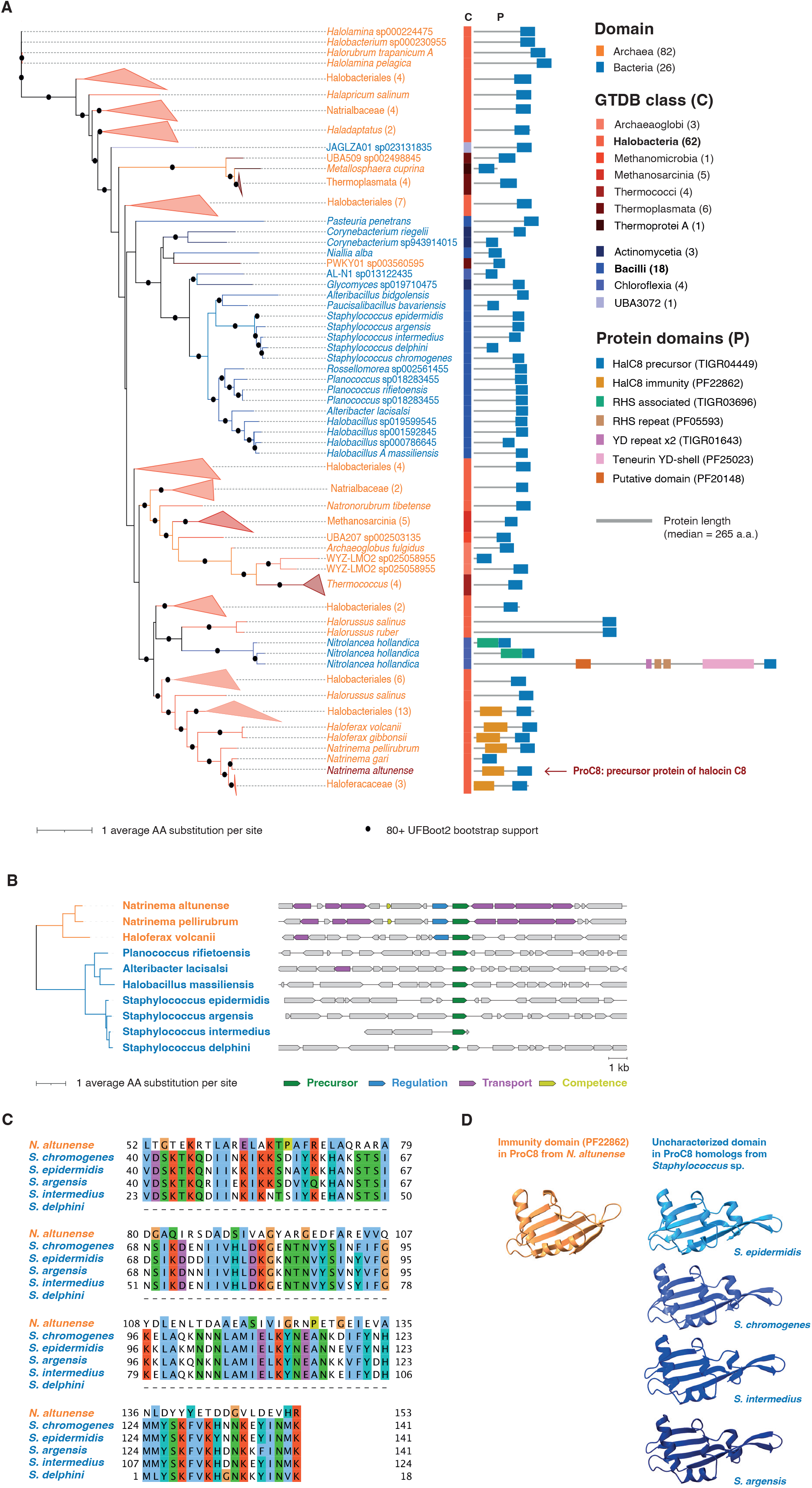
Halocin C8 and its bacterial homologs. **A.** Protein tree of halocin C8, highlighting domain composition and length of the respective homologs. **B.** Gene neighbourhood of halocin C8 homologs in archaeal and bacterial genomes. **C,D.** Sequence (C) and structural (D) alignment of the HalI immunity domain and the homologous regions in *Staphylococcus* spp.

A second, independent acquisition occurred in *Nitrolancea hollandica*, a nitrite-oxidizing bacterium (Sorokin et al. 2014) that appears to have acquired its C8 homologs – three in total – from *Halorussus* species. As previously described (Makarova et al. 2019), these proteins include rearrangement hot spot (RHS) repeats, RHS-associated domains and YD-repeats, which are features commonly found in polymorphic toxins. The presence of these features tentatively supports the hypothesis that halocin C8 in *N. hollandica* continues to be used in microbial conflicts.

Finally, a third acquisition of halocin C8 may have occurred in a member of bacterial phylum WOR-3.

Most intriguing to us is the presence, in the main group of bacteria, of five *Staphylococcus* genomes out of 77 in our database (6.5%), including *S. epidermidis*, a ubiquitous member of the human skin microbiome known to cause opportunistic infections (Otto 2009), *S. intermedius*, which is found on the skin of pigeons, cats, and dogs, and *S. delphini*, originally isolated from skin lesions of dolphins, but also recovered from the skin of other animals including foxes, minks and badgers (Varaldo et al. 1988; Guardabassi et al. 2012). It is interesting to note in this regard that haloarchaea have repeatedly been isolated from mammalian skin microbiomes although it remains unclear whether they stably inhabit these environments or represent contaminants, for example associated with the consumption of salty foods (Borrel et al. 2020; Umbach et al. 2021).

Gene expression data for *S. epidermidis* strain RP62A shows that its halocin C8-like protein is transcribed in stationary phase monocultures (Balasubramanian et al. 2018) and harbours a recognizable Sec signal peptide.

None of the proteins encoded in the neighbourhood of *halC8* have been characterized, so it remains unknown if they contribute in any way to C8 biology. We find no conservation of genes in the vicinity of C8 homologs in the *Staphylococcus* genomes of interest (Figure 4B), suggesting that C8, at least in these organisms, does not operate as part of a co-inherited and physically contiguous gene cluster.

HalI, the immunity protein that protects the original producer from halocin C8 and which is cleaved off from the precursor protein ProC8, is not found in bacteria when using the model and detection thresholds defined in Pfam (domain Hac8_like_N; PF22862). However, many bacterial C8 homologs have an N-terminal region of similar size to the original C8 (Figure 4A). We therefore decided to compare the structural features of the immunity domain in *N. altunense* with the AF3-predicted structure of the N-terminal region of *Staphylococcus* C8 proteins. We find that, despite divergent amino acid sequences (Figure 4C), the predicted 3D structure of the bacterial domains is striking similarity to that of the precursor domain of HalI (Figure 4D).

Does conservation of the immunity domain mean that some bacteria, including *Staphylococcus* species, may be susceptible to halocin C8? While Li and colleagues tested the activity of *Haloferax mediterranei* C8 on a handful of bacteria and reported no inhibition (Li et al. 2003) it is entirely possible that activity may be narrow and could affect certain bacteria exclusively. The structural conservation of HalI is intriguing and cautiously points to inter-bacterial conflicts.

## DISCUSSION

Archaea and bacteria can exchange genetic material. In certain environments, this exchange is extensive (Aravind et al. 1998; Zhaxybayeva et al. 2009). The results reported here, along with previous work (Makarova et al. 2019), suggest that systems used in intermicrobial conflict are no exception. Bacteriocins can be found in archaeal genomes, and archaeocins in bacterial genomes.

In the literature, we predominantly find use of bacteriocins by bacteria against other bacteria and use of archaeocins by archaea against other archaea. This might lead us to conclude that a bacteriocin, if present in an archaeal genome, will also be deployed against bacteria and that archaeocins in bacterial genomes will target archaea. However, the pre-eminence of within-Domain killing in the literature almost certainly reflects pervasive ascertainment bias. Bacteriocins simply do not get tested against archaea. It is entirely possible that systematic testing would reveal that some bacteriocins/archaeocins have broad-spectrum activity against both bacteria *and* archaea, as has already been demonstrated for halocin H4 (Chen et al., 2024). It is also possible that some bacteria-encoded bacteriocins only kill bacteria, but their archaeal homologs have evolved to kill archaea (and vice versa).

By computational analysis alone, we can neither conclusively establish that these cross-Domain acquisitions have retained their original function as tools of inter-microbial conflict nor whether bacteria or archaea (or both) are the principal targets of antagonistic activity. Ultimately, experimental investigation will be required to demonstrate continued use in conflict in general, and use in cross-Domain conflict in particular, but recent discoveries of bactericidal peptidoglycan hydrolases and phospholipases in archaea (Strock et al. 2025; Taissir et al. 2026) strengthen the view that archaea encode significant antibacterial capabilities that largely remain undescribed.

## METHODS

### Database of bacteriocins

Our starting point for building a database of bacteriocin core sequences is BAGEL4 (Heel et al. 2018), which provides amino acid sequences and hidden Markov models (HMMs) for a large number of known bacteriocins. However, this set contains many duplicated or very similar entries, so we first clustered the database of sequences using CD-HIT (Fu et al. 2012) with option ‘-c 0.9’. The final, de-duplicated set contains 423 sequences of proteins or precursor peptides from bacteriocin gene clusters.

### Bacteriocin search

For bacteriocins defined by HMM models, we searched our database of prokaryotic genomes (Strock et al. 2025) using the *hmmsearch* module of HMMER (hmmer.org), using the gathering threshold when available (‘--cut_ga’ option) or otherwise the bit score threshold of 20 (‘-T 20’ option). For bacteriocins defined by their amino acid sequence, we used the *phmmer* module with the same bit score threshold. This search yielded 10,904 protein hits (327 in archaea, 10,577 In bacteria). The 327 archaeal hits come from 85 unique bacteriocins (Table S1).

We then reduced the set of candidate bacteriocins in archaea by shortlisting only bacteriocins encoded in at least two archaeal genomes and removing hits for which no antibacterial activity has been experimentally demonstrated (comX homologs, Linocin M18, FlvA2f, FlvA2h, Piricyclamide 7005E2, SSV 2083, Caulonodin III, Ancovenin). One hit, Enterocin NKR 5 3D, had been observed to produce very weak activity, unlike the main peptide Enterocin NKR 5 3B (Ishibashi et al. 2012), which is not found in archaea, and was therefore removed. Similarly, we removed both Enterocin 1071A and Enterocin 1071B as both are required for antimicrobial activity (Balla et al. 2000) but only B is found in archaea. The final set of 15 bacteriocins all have experimental evidence of antibacterial activity.

### Genomes encoding subtilosin-like pathways

The precursor peptide of subtilosin A is encoded in RefSeq genome GCF_000009045.1 from *Bacillus subtilis* subsp. *subtilis* str. 168 (protein ID: NP_391616.1). Homologs in archaea were found in *Thermococcus celer* str. Vu 13 (RefSeq: GCF_002214365.1; protein ID: WP_088863144.1) and *Thermococcus nautili* str. 30-1 (RefSeq: GCF_000585495.1; protein ID: WP_042691590.1).

The amino acid sequences encoded by genes sboA, albA and albC and their homologs in *T. celer* and *T. nautili* were used to find other bacteria encoding a similar pathway. These sequences were searched against our database of prokaryotes using MMseqs2 (Steinegger and Söding 2017) with the highest sensitivity threshold (‘-s 7.5’ option). Genome hits with at least one of each three query genes, on the same contig and in a 20 kb region were kept. This resulted in the inclusion of eight additional genomes (*Bacillus spizizenii*, RefSeq: GCF_000227465.1; *Bacillus halotolerans*, RefSeq: GCF_001517105.1; *Bacillus atrophaeus,* RefSeq: GCF_001584335.1; *Bacillus smithii*, RefSeq: GCF_001050115.1; *Quasibacillus thermotolerans*, RefSeq: GCF_000812025.2; *Enterococcus devriesei*, RefSeq: GCF_001885905.1; *Aneurinibacillus thermoaerophilus*, RefSeq: GCF_900099925.1; *Caldicellulosiruptor danielii*, RefSeq: GCF_000955725.1).

### Phylogeny of sboA, albA and albC homologs

A single protein tree was built by aligning the amino acid sequences of the sboA, albA and albC homologs independently using MAFFT with the L-INS-I strategy (Katoh and Standley 2013), trimming the alignment using ClipKIT with the default ‘smart gap’ feature (Steenwyk et al. 2020), and concatenating the results. A maximum likelihood tree was then built with IQ-TREE 2 (Minh et al. 2020), using the Model Finder Plus algorithm (Kalyaanamoorthy et al. 2017) to automatically find a suitable substitution model (option ‘-m MFP’; the algorithm settled on ‘LG+R3’). Bootstrap value approximations were computed with UFBoot2 (Hoang et al. 2018) (option ‘-B 1000’). The tree was rendered with iTOL (Letunic and Bork 2024).

### Analysis of the subtilosin-like biosynthetic gene clusters

A window of 8 kb was considered downstream of the *sboA* gene homologs in all genomes encoding a putative pathway. Proteins encoded by genes in those windows were extracted and an all versus all search carried out using *phmmer* from the HMMER package to identify genes with significant homology to one of the original gene in the *sbo-alb* cluster. In addition, we identified a protein conserved in *T. celer* (protein ID: WP_088863146.1), *T. nautili* (WP_042691585.1) and *Caldicellulosiruptor danielii* (WP_045174440.1) as described above.

Sequence alignments were rendered with AliView (Larsson 2014). Structural alignments were computed and rendered with ChimeraX (Pettersen et al. 2021). Gene neighbourhoods were rendered with the Python library DnaFeaturesViewer (github.com/Edinburgh-Genome-Foundry/DnaFeaturesViewer).

### Halocin search

We considered three halocins with known precursor sequences: halocin H4, halocin C8 and halocin S8. The search was carried out on the entire GTDB release 214 as follows. For halocin C8, we first considered the protein domain ‘halocin_C8_dom’ (TIGR04449) from the TIGRFAMs dataset (Haft et al. 2001) and used the hmmsearch module from HMMER (with option ‘--cut_ga’ in order to use the gathering threshold defined in the model specification) to search our own prokaryotic database. This first pass provided a set of proteins that we used to seed the wider GTDB search. We performed two iterative searches using MMSeqs2. From this set, we made sure that all hits included a recognisable halocin C8 domain (TIGR04449). For halocin H4 and S8, no Pfam or TIGRFAMs models exist, so we searched GTDB release 214 directly with MMSeqs2 (Steinegger and Söding 2017) using three iterative searches.

### Halocin phylogeny and gene neighbourhoods

We constructed a maximum likelihood phylogenetic tree for halocin H4 using IQ-TREE version 3 (Wong et al. 2026) with automatic model finder option ‘-m MFP’ (Kalyaanamoorthy et al. 2017), which selected model ‘Q.PFAM+F+I+G4’, (Minh et al., 2021). The tree was rendered with iTOL (Letunic and Bork 2024). Bootstrap estimations were produced with UFBoot2 (Hoang et al. 2018).

The halocin C8 tree was constructed in the exact same way. The automatic model search procedure settled on substitution model ‘Q.PFAM+R5’. Some archaeal clades were collapsed in the final rendered tree for visualisation purpose. All proteins in these collapsed groups have the same domain architecture and very similar sizes. A single representative protein was picked at random to illustrate the clade.

Gene neighbourhoods were constructed as described above. For halocin C8, not all 208 gene neighbourhoods could be rendered, so we selected a subset of relevant archaea (including the original producer) and bacteria (including halophilic members and all five *Staphylococcus* species). *Staphylococcus chromogenes* was excluded because the contig containing the halocin C8 homologous protein does not contain any other genes.

### Structural investigation of the immunity domain of halocin C8

The protein structures of halocin C8 homologs from *N. altunense*, *S. epidermidis*, *S. chromogenes*, *S. delphini*, *S. intermedius* and *S. argensis* were predicted with AlphaFold 3 (Abramson et al. 2024). Structures were aligned using ChimeraX and only sections with significant structural homology to the immunity domain of the original producer were kept. The amino acid sequences underlying the structural alignment were extracted and rendered with JalView using the Clustal X colouring scheme.

## CODE & DATA AVAILABILITY

Code and supporting datasets can be found at https://github.com/srom/baab.

## ACKNOWLEDGEMENTS

This work was made possible by funding from the UKRI Medical Research Council (MC- A658-5TY40) to TW.

## SUPPLEMENTARY TABLES

**Table S1.** Bacteriocin homologs in archaea

**Table S2.** Archaeocin homologs in bacteria

